# MiCK: a database of gut microbial genes linked with chemoresistance in cancer patients

**DOI:** 10.1101/2024.07.24.604921

**Authors:** Muhammad Shahzaib, Haseeb Manzoor, Hasnain Zubair, Farzana Jabeen, Masood Ur Rehman Kayani

**Affiliations:** Department of Sciences, School of Interdisciplinary Engineering & Sciences (SINES), National University of Sciences & Technology (NUST), Srinagar Highway, Sector H-12, Islamabad, Pakistan; Department of Computing, School of Electrical Engineering and Computer Science (SEECS) National University of Sciences & Technology (NUST), Srinagar Highway, Sector H-12, Islamabad, Pakistan

**Keywords:** Cancer, Chemoresistance, Gut microbiome, Chemotherapy, Database, Metagenomics

## Abstract

Cancer remains a global health challenge, with significant morbidity and mortality rates. In 2020, cancer caused nearly 10 million deaths, making it the second leading cause of death worldwide. However, the emergence of chemoresistance becomes a major hurdle in successfully treating patients. Human gut microbes have been recognized for their role in modulating drug efficacy through their metabolites, ultimately leading to chemoresistance. The available databases are currently limited to knowledge regarding the interactions between gut microbiome and drugs. However, a database containing the human gut microbial gene sequences, and their effect on the efficacy of chemotherapy for cancer patients has not yet been reported. To address this challenge, we present the Microbial Chemoresistance Knowledgebase (MiCK), a comprehensive database cataloging microbial gene sequences associated with chemoresistance cancers. MiCK contains 1.6 million sequences of 29 gene types linked to chemoresistance and drug metabolism, curated manually from recent literature and sequence databases. The database supports efficient data retrieval and analysis, providing a user-friendly web interface for sequence search and download functionalities. MiCK aims to facilitate the understanding and mitigation of chemoresistance in cancers by serving as a valuable resource for researchers.

**Database URL:** https://microbialchemreskb.com/

## Introduction

Cancer is a significant global health concern, ranking among the leading causes of morbidity and mortality worldwide. According to the WHO, cancer accounted for nearly 10 million deaths in 2020, making it the second leading cause of death globally. The GLOBOCAN report highlights that there were approximately 19.3 million new cancer cases and 10 million cancer-related deaths in 2020 [1]. Factors contributing to the incidence of cancer include genetic predisposition, lifestyle factors such as diet, obesity, and alcohol consumption, as well as environmental exposures and infections. Multiple treatment strategies are available for cancer, ranging from surgical interventions to chemotherapy, radiation therapy, immunotherapy, and targeted therapies. The approach to treatment often depends on the type and stage of cancer, as well as the patient’s overall health and preferences [2].

Major drugs used to treat cancer include 5-Flurouracil (5-FU), Capecitabine, Gemcitabine, Irinotecan, and Oxaliplatin. 5-FU is a commonly used antimetabolite drug for cancer that causes the inhibition of thymidylate synthase, an enzyme essential for DNA synthesis and repair. Capecitabine, a prodrug of 5-FU, is converted to 5-FU in tumor tissue by the enzyme thymidine phosphorylase [3]. Furthermore, it has replaced 5-FU due to its improved pharmacokinetics and tolerability [4]. Gemcitabine is a nucleoside analog that inhibits DNA synthesis by interfering with the activity of the ribonucleotide reductase enzyme [5]. Irinotecan, a DNA topoisomerase inhibitor, binds to topoisomerase which causes the inhibition of DNA replication and transcription through its active metabolite i.e., SN38 [3]. Oxaliplatin works by forming DNA adducts inhibiting DNA replication, transcription, DNA damage, and apoptosis of cancer cells [3].

Chemoresistance is a major challenge in the successful treatment of cancer patients. The human gut microbiome has recently been attributed to the emergence of chemoresistance in cancer patients [7]. Gut microbial species including *E. coli, B. longum, Citrobacter*, and *E. faecalis* metabolize 5-FU, the most common drug used in treating cancer patients, and alter its efficacy [11]. Another prominent example includes the increased efflux of oxaliplatin by gut microbes through the glucuronidation mechanism by the β-glucuronidase enzyme. This enzyme inactivates SN38 (an activated form of irinotecan) by converting it to SN38G [10]. *F. nucleatum*, another common gut microbial species, promotes chemoresistance by the modulation of autophagy [8]. *Lactobacillus* and *Bifidobacterium* have been found to have higher abundance in the progressive disease group compared to the partial response group in cases administered with folinic acid, 5-FU, and irinotecan [6].

Several databases have been designed, considering the importance of the gut microbiome and its interaction with drugs used in chemotherapy. These include PharmacoMicrobiomic which contains 131 records about the effect of drugs on gut microbiome [7]. Microbe-Drug Association Database (MDAD) contains interactions of 1388 drugs and 180 microbes curated from drug databases and relevant literature [8]. MagMD is also a comprehensive database for metabolic actions of the gut microbiome on drugs, covering 32,678 interactions between 2,146 microbes, 36 enzymes, and 219 substrates [9]. To the best of our knowledge, no database currently exists that provides dedicated access to microbial gene sequences potentially involved in chemoresistance in cancer patients. To address this challenge, we present the Microbial Chemoresistance Knowledgebase (MiCK), a comprehensive database cataloging microbial gene sequences associated with chemoresistance in CRC. MiCK contains 1.6 million sequences of 29 gene types linked to chemoresistance and drug metabolism, curated from recent literature and databases. The database architecture, built using MySQL, supports efficient data retrieval and analysis, offering a user-friendly web interface for search and download functionalities. MiCK is designed to facilitate the understanding and mitigation of chemoresistance in cancer treatment, serving as an invaluable resource for researchers.

## Materials and Methods

### Identification of gut microbial chemoresistance genes

For the identification of gut microbial genes, potentially associated with chemoresistance, we first performed a literature search in PubMed [10] and Google Scholar [11] using the following keywords: “chemoresistance”, “cancer”, and “gut microbiome”. Through manual curation, the search results were restricted to only the publications that reported or discussed microbial genes or products that could potentially result in chemoresistance. In addition, enzymes involved in the metabolism of commonly used therapeutics for cancer were searched in the KEGG [12] and MagMD databases [9].

After collecting information on genes and enzymes we retrieved their sequences from the UniProtKB, and NCBI Gene databases [13]. Through this, we identified 29 gene types that could potentially confer chemoresistance. These gene types include Butyrate, short-chain fatty acid produced by gut microbes [14]; cytidine deaminase, produced by *Gammaproteobacteria* that inactivates Gemcitabine [15]; cysteine desulfurase (NFS1) weaken the sensitivity of cancer cells to oxaliplatin [16]; Dihydropyrimidine dehydrogenase (DHP), which causes catabolism of 5-FU [17]; Dutpase, whose expression is linked to susceptibility of cancer cells to chemotherapeutics like 5-FU [18]; Glutathione (GLU) [19]; Glutathione S-transferase (GST), a detoxification enzyme for oxaliplatin [20]; UDP-glucuronosyltransferases (UGTs) [21]; Histone deacetylase6 (HDAC6) [22]; NME2 (nucleoside diphosphate kinase 2) plays a role in chemoresistance to 5-FU, knockdown of NME2 gene increases sensitivity to 5-FU in CRC cell lines [23]; Ribonucleotide reductase consists of two subunits M1 and M2. M1 subunit is present in large amounts in microarray profiles of *in vivo* resistance to gemcitabine model which depicts its role in chemoresistance [24]; Thymidine kinases (TK) are involved in the salvage pathway capable of synthesizing deoxythymidine by phosphorylation of thymidine thus helping in DNA synthesis; TK2 involved in chemoresistance thus knocking them by small interfering RNA sensitizes the cell to gemcitabine chemotherapy [25]; 5-FU inhibits the thymidylate synthase (TS) stopping DNA replication and apoptosis of tumor cells. One of the key indicators of chemoresistance to 5-FU is an increase in the concentration of TS in tumor cells [19]; UMP/CMP kinase is an enzyme involve in the phosphorylation in 5-FU mechanism, but miR-130-b which is a key epigenetic regulator of UMP/CMP kinase attenuates the functioning and leads to chemoresistance of 5-FU [26]; β-glucuronidase (BGU) is an enzyme produced by mostly by four gut bacterial phyla i.e., *Bacteroidetes, Firmicutes, Verrucomicrobia*, and *Proteobacteria*. BGU induces toxicity in response to irinotecan thus limiting the dose and affecting the efficacy of irinotecan [27]; Most of the chemotherapeutics are topoisomerase active substances but the topoisomerase inhibitors (DTI) are involved in the chemoresistance against these drugs. DTI includes ATP-binding cassette transporters (ABC), GSH, and HDA6 [28]; Deoxycytidine kinase (*dck*) is involved in the phosphorylation and the studies reported the role of *dck* in limiting the cytotoxic activity of gemcitabine [29]; Hydroxyglutarate has been shown to inhibit α-ketoglutarate-dependent dioxygenases, including histone demethylases. This leads to epigenetic changes that can promote tumor progression and chemotherapeutic resistance [30]; Metallothionein (MT) are small cystine-enriched molecules with 4 isoforms, MT1 to MT4. Thiol groups of MT are involved in chemoresistance to oxaliplatin, a drug used for the treatment of CRC through the mechanism of drug detoxification which prevents cells from apoptosis and accumulation of oxaliplatin [31]; Toll-like receptors (TLR) are the main receptor pathway that can mitigate the bacterial process of production of pro-inflammatory cytokines that help in tumorigenesis and can lead to [32]chemoresistance in cancer cells. An active TLR-4-MyD88 signaling pathway could pose a risk for cancer development and serve as a promising target for creating biomodulators to combat chemoresistance [33]; Thymidine phosphorylase is involved in the activation of 5-FU, but it is also involved in tumor progression through angiogenesis and by avoiding apoptosis which depicts its role in chemoresistance against 5-FU based chemotherapy of cancer [34].

The gut microbiome can influence the functionality of drugs by secreting several enzymes [32]. Therefore, to enhance the pool of genes that can potentially result in chemoresistance, we searched for pathways of drug metabolism in the KEGG Pathway Database [12] and explored the MagMD database that provided the information about interaction of microbial enzymes and drugs [9]. We then retrieved the sequences of all the enzymes from UniProtKB and NCBI gene databases. β-ureidopropionase, phosphoribosyl transferase, uridine phosphorylase, uracil phosphoribosyl transferase, carboxyl esterase, uridine kinase, and deoxycytidine kinase were the enzymes involved in the metabolism of cancer drugs and were used in our database. The list of chemoresistance genes (CRGs) is provided in Table 1.

**Table 1:**
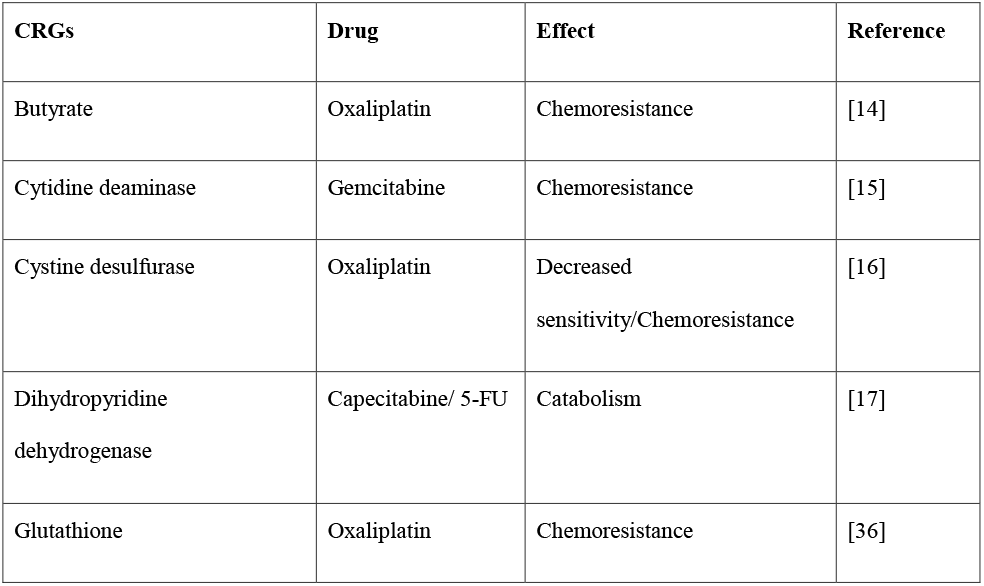
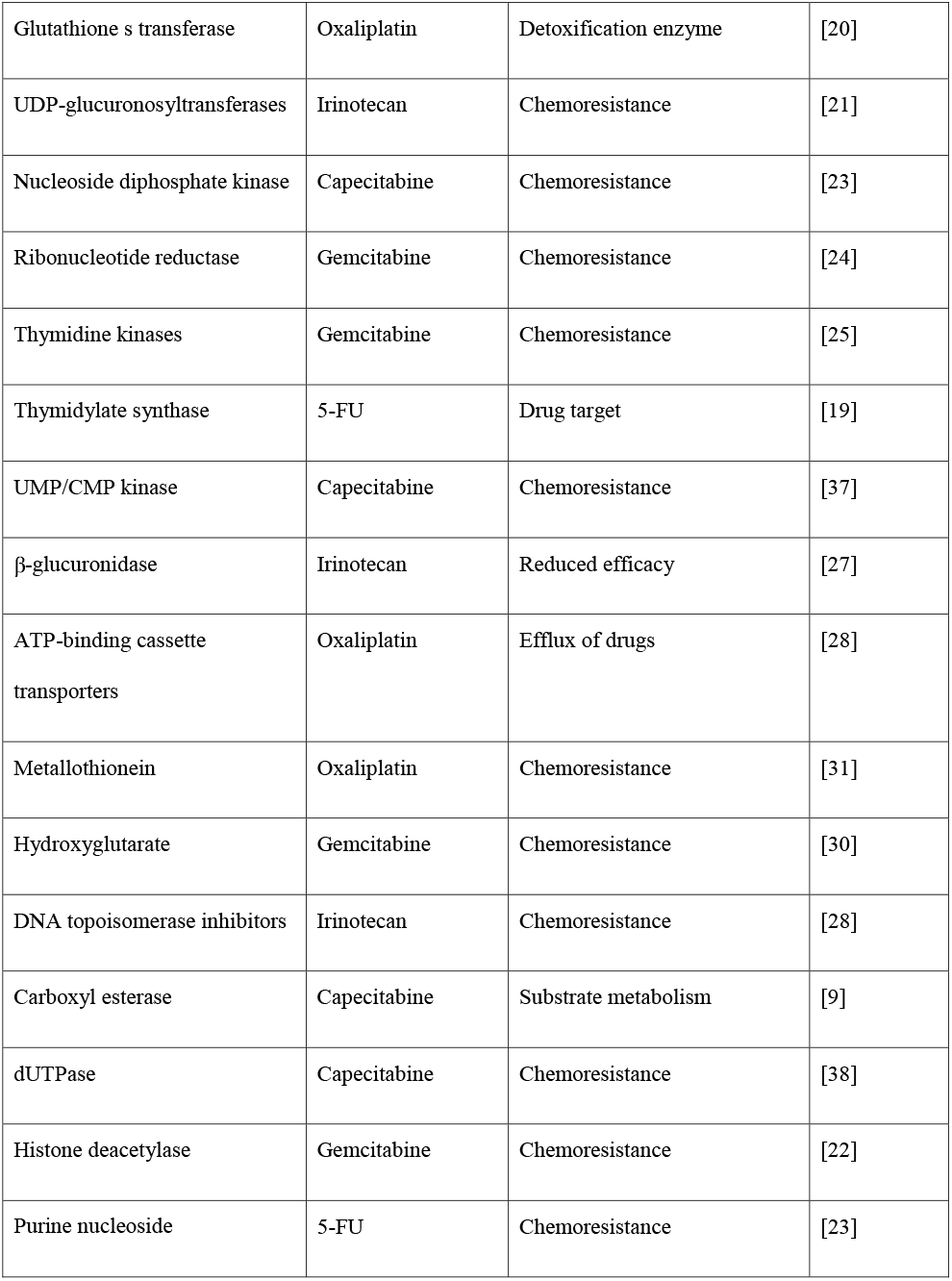
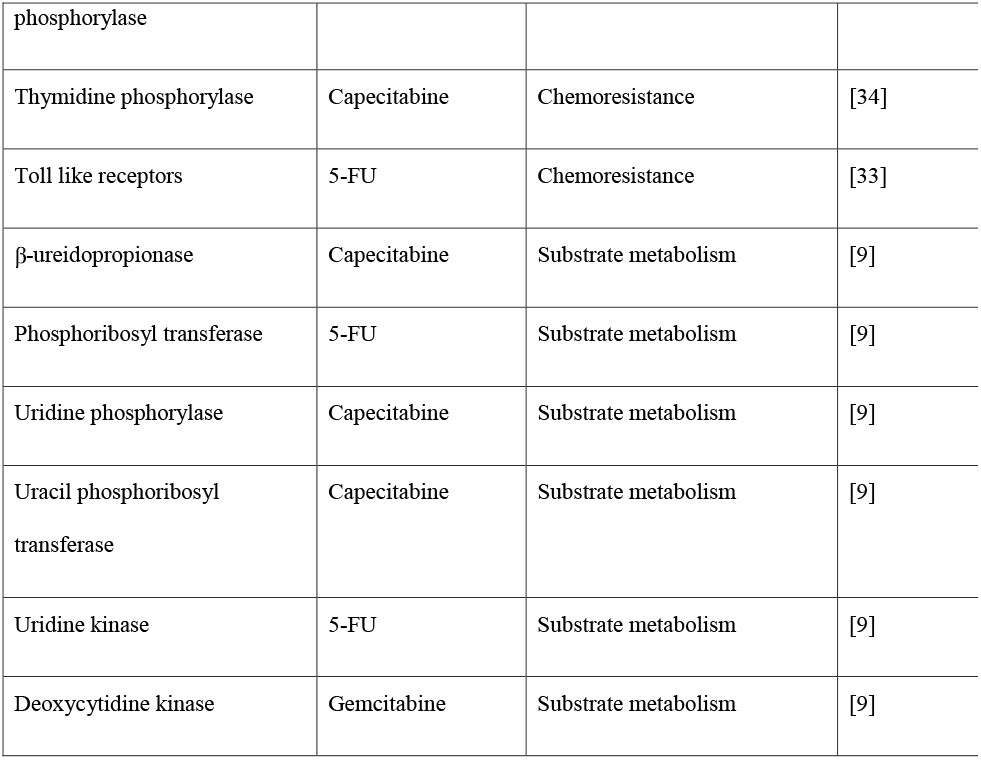
Chemoresistance genes, associated drugs, and their effects.

### Non-redundant CRG catalogue and database construction

MMseqs2 (version 13.45111) was used to cluster the CRGs [35]. MMseqs2 used a clustering algorithm to construct the indexed database from the concatenated sequences file and then performed clustering using linclust. Clustering was performed using multiple identity thresholds (i.e., 50%, 75%, 80%, 90%, and 100%) with query coverage of 80%.

The database was created using MySQL (version 8.0.2, https://www.mysql.com/) consists of 5 columns: “Accession_number”, “Gene_type”, “Drug”, “Uniport_Accession”, and “Effects”. The accession number serves as the primary key of the database, the second column contains gene type information, the third column provides the target drug for the gene type, the fourth column holds the full gene type names, the fifth column contains the original accession numbers for retrieving sequences from UniProtKB, and the last column holds the information of effects on the drugs. The framework of the database construction is illustrated in Figure 1.

**Figure 1:**
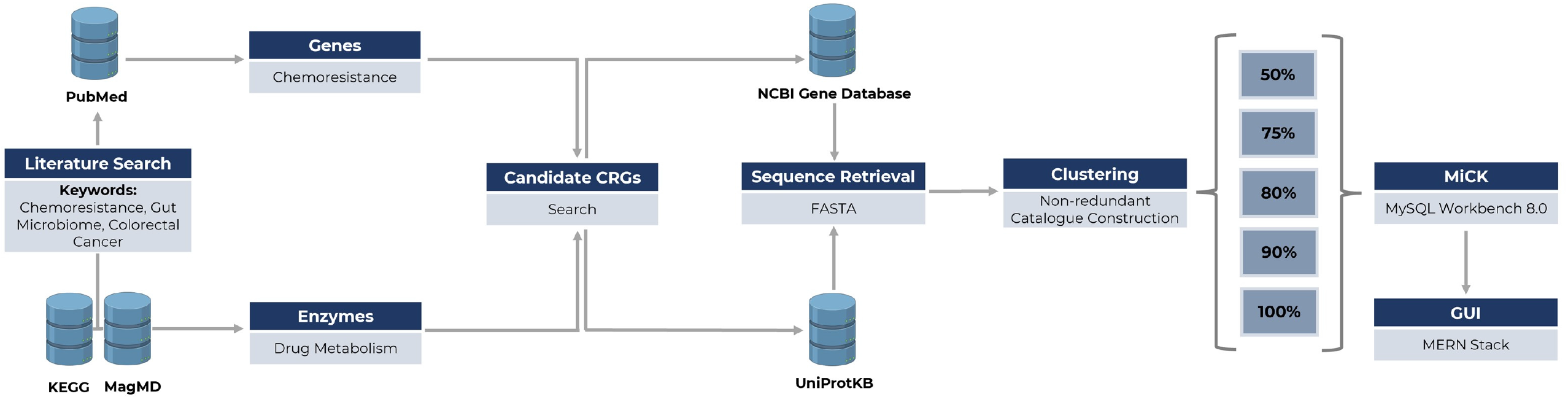
Framework for developing MiCK.

### Graphical user interface design

The MiCK platform is developed using the MERN stack, which includes MongoDB, Express, React, and Node.js. MongoDB was utilized as the database. Express.js served as the backend framework to communicate between the frontend and database. React.js was employed for building the user interface, offering a responsive and interactive environment for researchers to input, view, and analyze data.

### MiCK Statistics

MiCK hosts ∼1.6 million sequences that encompass 29 CRGs following a thorough literature search and an extensive review of drug-metabolizing enzymes. Among these GLU is the most frequent gene type included in MiCK with ∼0.6 million representative gene sequences, followed by PRT (∼0.25 million gene sequences) (Figure 2A). MiCK contains ∼0.6 million genes that target oxaliplatin, followed by gemcitabine for which ∼0.4 million target gene sequences are present in MiCK (Figure 2B). The effects column comprises chemoresistance, chemosensitivity, and substrate metabolism. Chemoresistance is the most frequent effect with almost 1.3 million repetitions.

**Figure 2:**
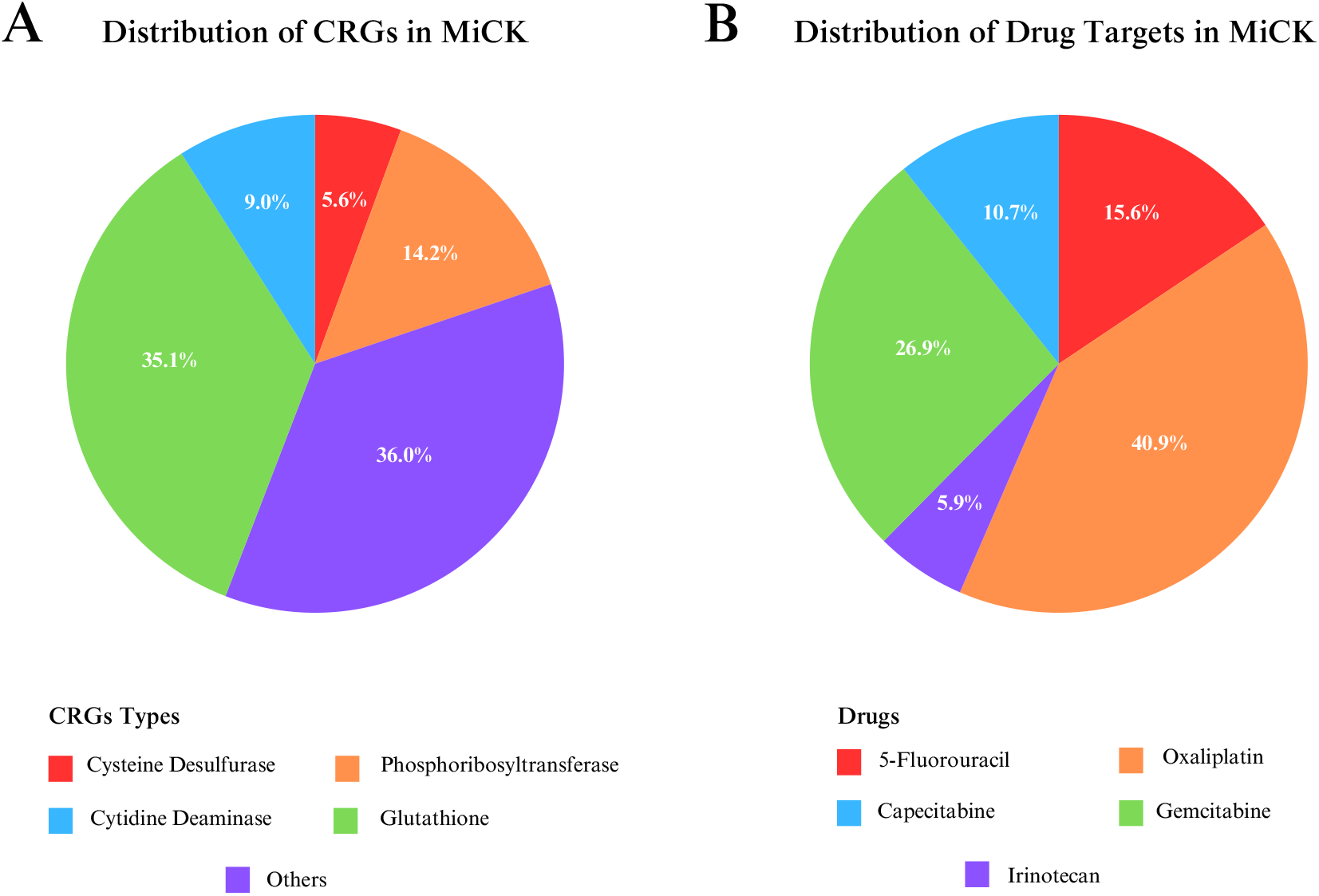
Key statistics of MiCK. (A) Distribution of chemoresistance genes in the MiCK. (B) Distribution of Drug Targets in the MiCK database.

### Database web interface

MiCK offers a user-friendly interface with 4 navigation bars, i.e. Home, Search, Download, and About.

#### 1. Home

The home page serves as an introductory page comprising of general information about the database. It serves as an informative desk for other pages, i.e., how to navigate, search and download through MiCK. The home page also presents users with the key statistics with visualization of pie charts.

#### 2. Search

The search page comprises a selection box, where you select the search criteria; search box, that allows keyword input; and a search button to initiate the search process. The search page allows a keyword search across the database. Users can search through accession number, drug name, gene name, effect, or bacteria name. The resultant table displays the information of gene type, its drug target, potential effect on the drug, and the associated bacterial host.

#### 3. Download

The download page offers the utility to download the CRG catalog which includes the sequencing files as well as the metadata. Currently, users can download the CRG catalog clustered at 50%, 75%, 80%, 90%, and 100% similarity cutoffs. Metadata includes information about the CRGs, corresponding drugs, effects on the drug. The graphical user interface of MiCK is shown in Figure 3.

**Figure 3:**
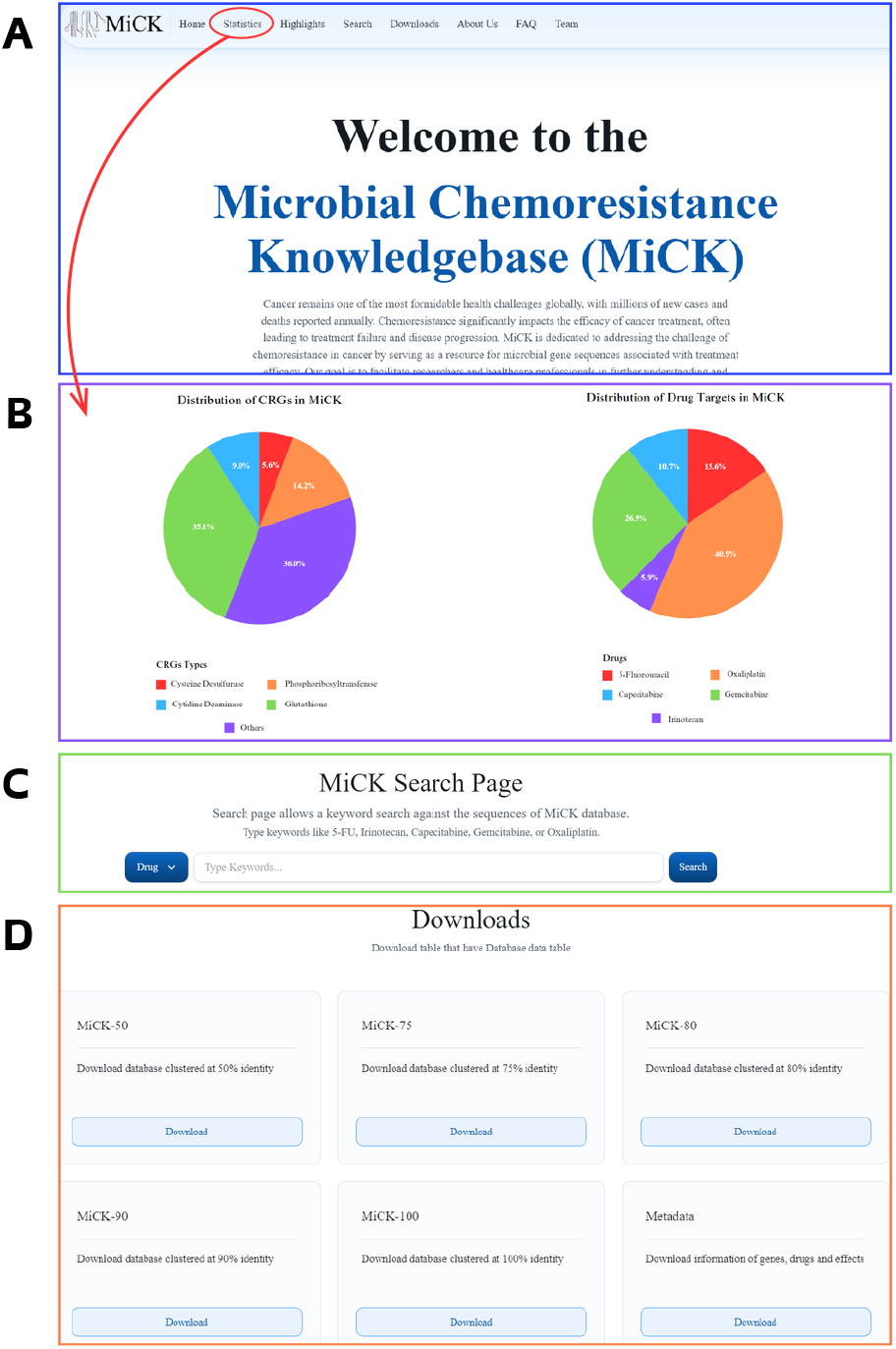
The graphical user interface of MiCK. (A) The Home page of MiCK. (B) The key statistics of MiCK. (C) The Search page of MiCK. The user can search gut microbial genes and their effect on drugs. (D) The Download page provides a utility to download datasets used in MiCK.

#### 4. About

The About page provides help for the users by answering some of the most frequently asked questions (FAQs) that a potential user can have in mind. In addition, the contact information of the database administrator is also provided for further queries.

## List of abbreviations

Abbreviation: Full form
UniProtKB: Universal Protein Knowledgebase
5-FU: 5-Fluorouracil
CRC: Colorectal cancer
GLOBOCAN: Global Cancer Observatory
KEGG Kyoto: Encyclopedia of Genes and Genomes
MDR: Multidrug resistance
RT-PCR: Reverse transcription polymerase chain reaction
SCFA: Short-chain fatty acids
CRG: Chemoresistance genes

## Declarations

### Availability of data and material

The publicly retrieved gene sequences and relevant details are provided in Supplementary Table 1.

### Author’s contributions

MS performed the literature search, curation, and sequence retrieval. HM, and HZ performed the database construction. FJ developed the GUI for the database. MRK conceived the idea, supervised the research, and provided resources for the project. All authors contributed to manuscript preparation and have read and agreed to the submission of the manuscript.

## Acknowledgments

The authors are highly thankful to Umme Habiba, M. Faheem Raziq, M. Adnan Tariq, and members of the Microbiome Research Group at the SINES, NUST, Islamabad, Pakistan, for useful discussion and feedback on our manuscript.

